# Specific targeting of plasmids with Argonaute enables genome editing

**DOI:** 10.1101/2022.04.14.488398

**Authors:** Daria Esyunina, Anastasiia Okhtienko, Anna Olina, Maria Prostova, Alexei A. Aravin, Andrey Kulbachinskiy

## Abstract

Prokaryotic Argonautes (pAgos) are programmable nucleases involved in cell defense against invading DNA. Recent studies showed that pAgos can bind small single-stranded guide DNAs (gDNAs) to recognize and cleave complementary DNA *in vitro. In vivo* pAgos preferentially target plasmids, phages and multicopy genetic elements. Here, we reveal that CbAgo nuclease from *Clostridium butyricum* can be used for genomic DNA cleavage and engineering in bacteria. CbAgo-dependent targeting of genomic loci with plasmid-derived gDNAs promotes recombination between plasmid and chromosomal DNA. Efficient genome cleavage and recombineering depends on the catalytic activity of CbAgo, its interactions with gDNAs, and the extent of homology between plasmid and chromosomal sequences. Specific targeting of plasmids with Argonautes can be used to integrate plasmid-encoded sequences into the chromosome thus enabling genome editing.

**One-Sentence Summary:** Prokaryotic Argonaute nuclease induces DNA interference between plasmid and chromosomal DNA to promote genome recombineering.

## Main Text

Prokaryotic Argonautes proteins (pAgos) are ancestors of eukaryotic Argonautes (eAgos), which act as central components of eukaryotic RNA interference and are involved in silencing of foreign elements and gene regulation (*1-6*). Intriguingly, while eAgos bind small RNA guides to recognize and silence RNA targets, all known pAgos preferentially target DNA (*6-8*). Nucleic acid cleavage by Argonautes depends on an RNaseH-like nuclease site in the PIWI domain, with the site of cleavage located precisely between the 10^th^ and 11^th^ guide nucleotides. Analysis of pAgos from several thermophilic and mesophilic prokaryotes demonstrated that they can be programmed with small (∼18 nt) DNA or RNA guides to cleave single-stranded DNA with high specificity (*9-18*). However, their activity toward double-stranded DNA *in vitro* is limited due to their inability to unwind DNA strands (*11, 13, 16*), and cellular functions of pAgos have remained largely unknown. Recently, CbAgo from *C. butyricum* was shown to defend heterologous *Escherichia coli* host from bacteriophage infection; several pAgos including CbAgo were found to preferentially target plasmid DNA suggesting that they function in cell defense against invaders (*19*).

In bacterial cells, CbAgo is primarily loaded with small gDNAs of plasmid origin and can use them to recognize and cleave homologous genomic loci and neighbor regions in the process of DNA interference (*19*). CbAgo also attacks naturally occuring or engineered double-strand breaks (DSBs) in cellular DNA (*19*). Preferential targeting of plasmids, bacteriophages and other multicopy elements by pAgos is likely explained by their intense replication, high copy numbers and high frequency of DSBs, resulting in their efficient processing by a combined action of pAgo and the cellular helicase-nuclease RecBCD (*19*). RecBCD rapidly degrades foreign DNA but promotes repair of genomic DNA through homologous recombination due to the presence of Chi-sites in the host genome, which switch the RecBCD activity from DSB processing to RecA loading onto the 3’-ended DNA strands (*20, 21*). The distribution of gDNAs bound by CbAgo at its target regions in genomic DNA is dependent on Chi-sites indicating the involvement of RecBCD in gDNA generation during DSB processing (*19*).

Here, we sought to elucidate the requirements for genomic DNA cleavage by CbAgo and to reveal whether it can be used for genome engineering. We used CbAgo to introduce targeted DNA breaks at specific loci of the bacterial chromosome and showed that it can induce highly efficient integration of plasmid-encoded sequences into the homologous chromosomal locus.

### CbAgo targets bacterial chromosome during DNA interference

To determine the minimal length of homology region required to induce DNA interference between plasmid and genomic DNA, we cloned fragments of the chromosomal *lacI* gene into the same plasmid that was used for expression of CbAgo (50, 100, 200, 300, 450, 600 bp or full-length 1083 bp *lacI*). In addition, the plasmid contained the full-length *araC* gene (876 bp) also present in the chromosome. We transformed *E. coli* with these plasmids, induced expression of CbAgo, purified the protein, and isolated and sequenced associated gDNAs (Fig. 1A). Their mapping to the plasmid and genome sequences showed that CbAgo preferentially binds plasmid gDNAs, as previously reported (*19*). After accounting for the plasmid copy number, ∼3.5-5 fold enrichment of plasmid-derived gDNAs was observed in all small DNA libraries (Table S1).

**Fig. 1.**
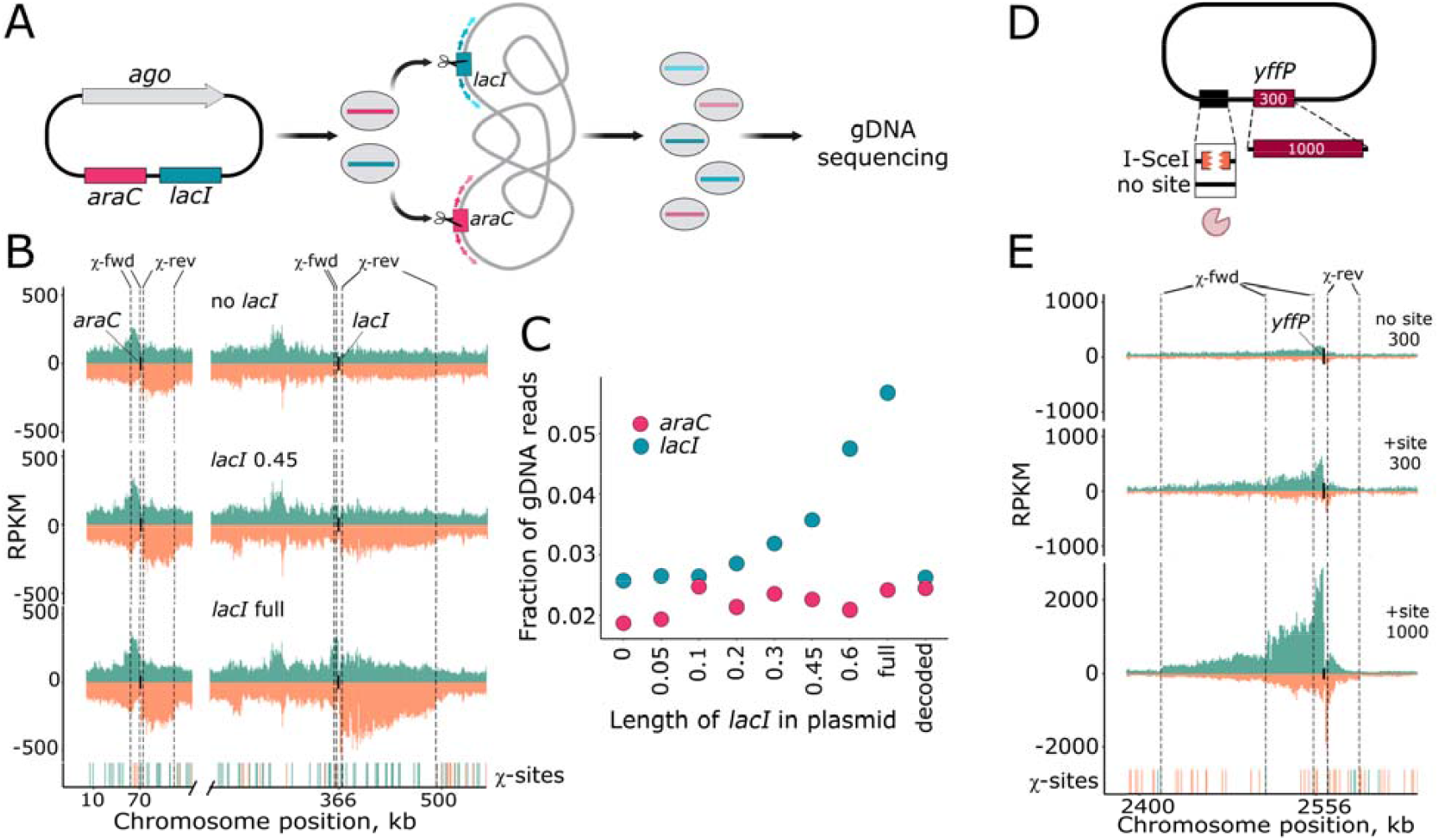
Requirements for genomic DNA cleavage by CbAgo. (A) Scheme of the DNA interference assay. Plasmid encoding CbAgo, *araC* and fragments of the *lacI* gene of various lengths is used by CbAgo as a source of gDNAs to attack homologous genomic regions, followed by their intense processing and loading of new gDNAs into CbAgo. (B) Analysis of gDNAs associated with CbAgo in *E. coli* strains containing plasmids with fragments of the *lacI* gene. Strand-specific distribution of gDNAs around the *araC* (left) and *lacI* (right) genes is shown for no *lacI*, 0.45 kb *lacI* and full-length *lacI* libraries; reads from the plus and minus genomic strands are shown in green and orange, respectively. The closest Chi sites surrounding the target genes in the proper orientation are indicated (forward for the plus strand and reverse for the minus strand). (C) The fraction of gDNAs corresponding to the *araC* (pink) and *lacI* (green) loci depending on the length of the plasmid *lacI* fragment. Fraction of gDNA reads between the four innermost Chi-sites around *araC* or *lacI* (indicated with dotted lines in panel B) was calculated relative to the total number of gDNAs mapped to the *E. colli* chromosome. (D) Scheme of the plasmid with an engineered I-SceI cut site and a single genome homology region, *yffP* gene (300 or 1000 bp). (E) Analysis of gDNAs associated with CbAgo in *E. coli* strains depending on DSB in plasmid DNA. Strand-specific distribution of gDNAs around the *yffP* gene is shown (reads from the plus and minus genomic strands are shown in green and orange, respectively). Several closest Chi sites surrounding the target genes in the proper orientation are indicated (forward for the plus strand and reverse for the minus strand).

Analysis of the chromosomal distribution of gDNAs revealed their enrichment at the sites of replication termination, *terA* and *terC* (Fig. S1A), as a result of intense processing of genomic DNA by RecBCD in this region (*19*). Similarly, SeAgo and TtAgo were previously shown to target the *ter* region in *Synechococcus elongatus* and *Thermus thermophilus* (*14, 22*).

The highest gDNA peaks outside of the *ter* region were observed around the *lacI* and *araC* genes, as a result of genomic DNA processing in these regions triggered by DSBs introduced at *lacI* and *araC* by CbAgo loaded with plasmid gDNAs corresponding to these genes (Fig. S1A). The size of the *araC* peak was constant in all analyzed strains (Fig. 1B,C and Fig. S2). In contrast, the *lacI* peak was absent in the case of plasmids with *lacI* fragments < 300 bp or lacking *lacI*, was fairly visible with a 300 bp *lacI* fragment, and was gradually increased in the case of plasmids with 450 bp, 600 bp and full-length *lacI* (Fig. 1B,C and Fig. S2). This indicates that >300 bp region of homology is sufficient to induce genomic DNA cleavage during DNA interference, and that the efficiency of DNA interference directly correlates with the fragment length. Recoding of the *lacI* gene by introducing single-nucleotide substitutions in its every 6^th^-8^th^ position resulted in disappearance of the gDNA peak around genomic *lacI* indicating that high level of homology between plasmid and target genes is required to induce DNA interference (Fig. S1C, Fig. S2).

The gDNA peaks around *lacI* and *araC* were much wider than the target genes (including dozens or even hundred kilobases from both sides, Fig. 1B, Fig. S2), indicating that the targeted chromosomal region is extensively processed after introducing of DSBs with CbAgo. The distribution of gDNAs around the target genes is asymmetric, with most gDNAs generated from the 3’-ended DNA strands facing the target gene. On both sides, sharp drop in the amounts of gDNAs is observed at several (one-three) Chi-sites oriented toward the target gene (Fig. 1B, Fig. S2). This indicates that DSB processing and gDNA generation is performed by RecBCD together with CbAgo (*19*).

To check whether cleavage of genomic DNA by CbAgo might affect transcription of target genes, we analyzed their mRNA levels in *E. coli* strains with and without DNA interference. It was found that introduction of DSBs in the *lacI* and *araC* loci by CbAgo does not significantly change expression of the *lacI, lacZ* and *araC* genes (Fig. S3), indicating that these DSBs are likely rapidly repaired without affecting transcription.

Processing of plasmid DNA by RecBCD generates gDNAs bound by CbAgo (*19*) and may therefore enhance DNA interference. To test whether plasmid targeting by CbAgo can be stimulated by DSB formation, we introduced the recognition site of the I-SceI meganuclease in plasmids containing either 300 bp or 1000 bp fragments of the *E. coli yffP* operon (Fig. 1D). CbAgo and I-SceI were expressed from the chromosome to prevent changes in protein expression because of plasmid degradation. We isolated gDNAs bound to CbAgo in *E. coli* strains containing these plasmids and analyzed their genomic distribution. All three profiles looked similar, except for the *yffP* gene area (Fig. S1B). In the case of the plasmid with a 300 bp fragment of *yffP* in the absence of the I-SceI cut site, only a small peak of gDNAs was visible around *yffP* in chromosomal DNA (Fig. 1E), in agreement with the results obtained for *lacI* (see above). In the presence of the I-SceI site, the size of the peak was notably increased indicating that DSB in plasmid DNA enhances DNA interference. Furthermore, the peak was dramatically expanded in the presence of the I-SceI site and a 1000 bp fragment of the *yffP* operon, with its size being comparable with the peaks at *ter* sites (Fig. 1E, Fig. S1). Increased cleavage of chromosomal DNA in the presence of the plasmid I-SceI site correlated with increased loading of plasmid-derived gDNAs into CbAgo (16.7 and 16.9-fold enrichment for the plasmids containing 300 and 1000 bp fragments of the *yffP* operon, in comparison with 6.9-fold enrichment for the plasmid lacking the I-SceI site). This indicates that engineered DSB in plasmid DNA stimulates its processing and generation of gDNAs bound by CbAgo, further increasing cleavage of the homologous chromosomal locus.

### CbAgo stimulates homologous recombination

The rate of homologous recombination between plasmid and chromosomal DNA in bacteria is very low thus preventing direct screening of recombinants (*23*). The Red recombinase of phage λ was used to increase the frequency of recombination (*24, 25*) but it still requires additional steps to remove antibiotic resistance markers or eliminate non-edited cells for generation of marker-less mutations, *e*.*g*. by negative selection using CRISPR-Cas (*26, 27*). The ability of CbAgo to target plasmids and generate DSBs in chromosomal DNA using plasmid-derived gDNAs might provide a possibility for efficient genome recombineering using plasmids homologous to the target locus.

As a proof of concept, we analyzed recombination between the genomic *lac* locus and a thermosensitive editing plasmid containing this locus (Fig. 1B). The *lacI* gene in the middle of the plasmid homology region was replaced with an antibiotic resistance marker (chloramphenicol resistance, Cm^R^) to detect recombination events, and the surrounding regions contained natural Chi-sites to promote recombination. The editing plasmid also contained a second resistance gene (spectinomycin resistance, Sp^R^), which was located outside of the homology region (Fig. 2A). CbAgo was expressed from an arabinose inducible promoter from a second plasmid, which contained the *lacI* gene to direct CbAgo to the target genomic locus (the same plasmid was used in Fig. 1A-C). *E. coli* cells were transformed with both plasmids and grown at the permissible temperature for the editing plasmid, either in the presence of arabinose to induce expression of CbAgo or in the presence of glucose to repress it. After one passage at 30 °C, the cells were grown at 43 °C to cure the editing plasmid, and the percentage of Cm^R^, Sp^R^, and Cm^R^Sp^R^ cells was calculated. Sp^R^ cells and double resistant Cm^R^Sp^R^ cells correspond to bacteria that have not cured the plasmid and contain both resistance markers. In contrast, Cm^R^ cells that have lost spectinomycin resistance correspond to bacteria that have undergone recombination and integrated the Cm^R^ gene into the chromosome.

**Fig. 2.**
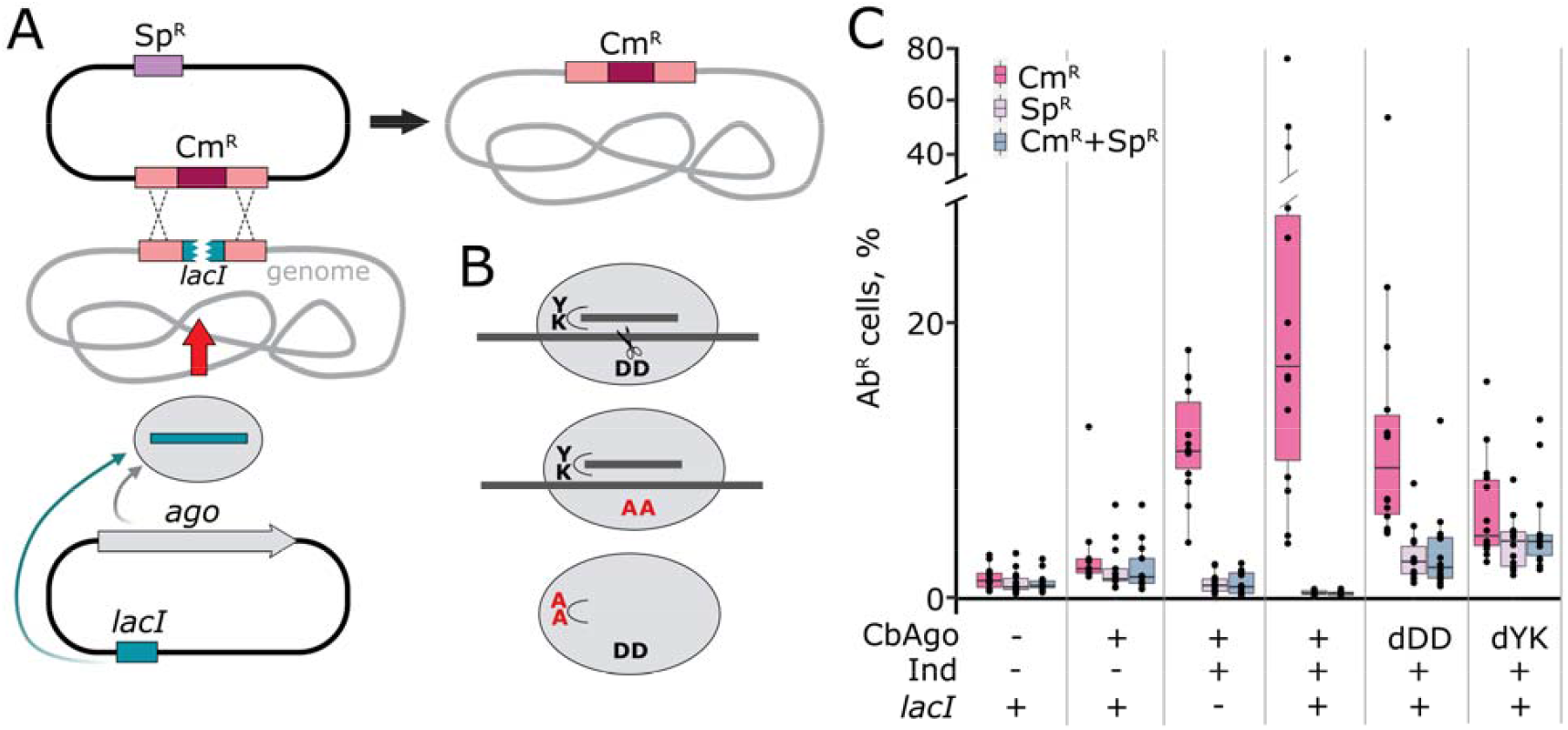
Genomic recombination induced by CbAgo. (A) Scheme of CbAgo-dependent recombination between homologous plasmid and chromosomal loci. The first plasmid encodes CbAgo and the *lacI* gene; the second plasmid contains a part of the *lacI* locus, in which *lacI* is completely replaced with the *cat* gene (Cm^R^), and the Sp^R^ gene in the plasmid backbone. After successful recombination, the *lacI* gene in the chromosome is replaced by the *cat* gene. (B) Mmutant variants of CbAgo: dDD CbAgo with alanine substitutions of two catalytic aspartates (DD>AA) that has no catalytiic activity, and dYK CbAgo (YK>AA) with impaired gDNA binding. (C) Frequencies of recombination in the target chromosomal locus. Percentage of Cm^R^, Sp^R^, and Sp^R^Cm^R^ *E. coli* cells after plasmid curing is indicated for each experimental condition. Dots represent data from independent biological replicates. Experimental conditions are indicated below the plot, including the presence of the CbAgo and *lacI* genes on the first plasmid, and CbAgo expression (either silenced with 1% Glc or induced with 0.1% Ara).

In all experimental conditions, the proportion of Sp^R^ and double Cm^R^Sp^R^ cells were similar and in the range of 0.3-4% indicating that plasmid was lost by the majority of cells (Fig. 2C). In the absence of the CbAgo gene or in the absence of CbAgo expression, the number of Cm^R^ cells (1.5-3%) were similar to the number of Sp^R^ and Cm^R^Sp^R^ cells, indicating that these cells harbor the remaining editing plasmid and are not recombinants. In contrast, when CbAgo expression was induced with arabinose, the percentage of Cm^R^ cells increased dramatically (up to 77% in some biological replicates, 23% on average), while the number of Sp^R^ and Cm^R^Sp^R^ cells that retained the editing plasmid significantly decreased (0.3% on average in comparison with 2% without CbAgo induction) (Fig. 2C, Table S2). Therefore, CbAgo not only allows genomic integration of the reporter cassette but also helps to eliminate the editing plasmid from the bacterial population. This results in ∼70-fold excess of cells with the genomic integration of the Cm^R^ gene over cells that have not lost the plasmid, in comparison with ∼1.5-fold difference without CbAgo expression. Importantly, the increase in the frequency of recombination in the presence of CbAgo may be likely higher since we cannot estimate the proportion of cells with the integrated Cm^R^ gene in control strains in the absence of CbAgo expression (and their numbers are likely much lower than the fraction of the cells that retain the editing plasmid).

To test whether the catalytic activity of CbAgo is required for efficient recombination, we performed a similar experiment with a mutant variant of CbAgo with alanine substitutions of two of its catalytic tetrad residues that disrupt its nuclease activity (dCbAgo, ‘dDD’) (Fig. 2B) (*13, 19*). With this mutant, the frequency of Cm^R^ cells was significantly decreased in comparison with active CbAgo (13% vs. 23%), while the proportion of cells that have not cured the plasmid increased (3-3.3% vs. 0.3%) (Fig. 2C). As a result, the ratio of Cm^R^ to Sp^R^ cells was strongly decreased (4.4-fold difference vs. 70-fold for wild-type CbAgo). At the same time, the fraction of Cm^R^ cells with dCbAgo was significantly higher than in the absence of CbAgo, indicating that dCbAgo can promote recombination even in the absence of nuclease activity. Previously, dCbAgo was shown to be loaded with gDNAs *in vivo* (*19*), indicating that it might still recognise the target chromosomal locus and promote homologous recombination without direct DNA cleavage (*e*.*g*., by generating extended single-strand DNA regions in this locus).

In contrast to dCbAgo, substitutions of two residues in the guide binding pocket in the MID domain in CbAgo (YK>AA, ‘dYK’, Fig. 2B) resulted in a much stronger defect of its recombineering activity. The substituted residues play a crucial role in guide DNA binding by both pAgos and eAgos (*16, 17, 28-30*). Accordingly, the recombination frequency measured with this mutant was strongly decreased in comparison with wild-type CbAgo while the efficiency of plasmid elimination was further increased (Fig. 2C). As a result, there was no significant difference in the number of Cm^R^ and Sp^R^ cells with this variant of CbAgo.

Since the editing plasmid already contains regions of homology to chromosomal DNA, we hypothesized that the presence of the *lacI* gene in the CbAgo plasmid may not be absolutely required for successful genome editing. Indeed, an increased frequency of recombination was also observed in the absence of *lacI* with wild-type CbAgo (11% of Cm^R^ cells on average in comparison with 1% of Sp^R^ cells that have not lost plasmid in the same population) (Fig. 2C, *lacI*-minus conditions).

In all previous experiments, chloramphenicol was added simultaneously with CbAgo expression, to minimize loss of the editing plasmid before recombination. To confirm that selection for chloramphenicol resistance is not required for efficient generation of genomic recombinants, we performed a similar experiment (using the plasmid with wild-type CbAgo and the *lacI* gene), but without the addition of chloramphenicol. The average frequency of Cm^R^ cells in this experiment was 9.3% (11.8, 2.4, 17.7 and 5.6% in four biological replicates), in comparison with 0.12% (0.25, 0.04, 0.1, 0.08%) of Sp^R^ cells (∼80-fold difference), indicating efficient genomic integration of the Cm^R^ gene.

Overall, these results demonstrate that CbAgo enables specfic integration of plasmid-encoded genes into the chromosome, and the high efficiency of recombination potentially allows obtaining of bacterial recombinants without antibiotic selection.

### DNA interference by pAgos enables genome editing

This and previous studies (*19*) demonstrated that CbAgo can induce DNA interference between homologous plasmid and chromosomal loci. Preferential targeting of plasmids was observed for several pAgos (*15, 16, 19*); using CbAgo, it was shown that this depends on the combined action of pAgo and the RecBCD nuclease-helicase, resulting in biased generation of gDNAs from plasmid DNA (*19*). Efficient loading of CbAgo with plasmid-derived gDNAs allows it to introduce DSBs in specific genomic regions, with the minimal length of homology region being in the order of half a kilobase. This size likely corresponds to the minimal length of an invading genetic element that can be targeted and eliminated by pAgos during DNA interference. Furthermore, the efficiency of plasmid processing and chromosome cleavage can be increased in the presence of engineered DSBs in plasmid DNA, even when using shorter homology regions, likely because DSBs increase production of gDNAs resulting in enhanced cleavage of the chromosome.

We demonstrate that CbAgo promotes integration of plasmid-encoded genes into the the target locus of the chromosome. We show that recombination depends on the catalytic activity of CbAgo and its interactions with gDNAs. The high efficiency of recombination with plasmid DNA is likely achieved by (1) continued DNA cleavage in the chromosomal target locus by pAgo loaded with plasmid gDNAs, (2) homologous recombination of the cleaved locus with the plasmid editing template, (3) elimination of plasmid DNA by pAgo.

Coordinated processing of plasmid and genomic DNA is likely promoted by cooperation of CbAgo with other nucleases involved in cell defense against invading DNA, including RecBCD (*19*). pAgos are also often encoded in prokaryotic genomes together with CRISPR-Cas suggesting that specific cleavage of foreign DNA sequences by CRISPR-Cas nucleases may similarly enhance their targeting and destroying by pAgos (*19*). At the same time, we hypothesize that Ago-mediated recombination could play a role in horizontal gene transfer in prokaryotes. Since there is a constant flow of DNA in the bacterial world via plasmids, phages and during natural transformation, induction of DNA interference after uptake of homologous sequences may lead to their efficient integration together with additional nonhomologous genes.

In comparison with other types of programmable nucleases, including TALEN, Zn-finger and Cas nucleases, pAgos do not require sophisticated protein engineering for target recognition and can be easily programmed with small DNAs without the requirement for specific motifs in the guide of target sequences. In contrast to CRISPR-Cas nucleases, pAgos do not require additional expression of specific guide RNAs and can be autonomously loaded with gDNAs generated from plasmid DNA, which allows their programming with the same plasmid sequences that participate in recombination. In wild-type *E. coli* cells, recombination requires the action of the RecBCD system. We envision that using pAgos in combination with the λ Red recombinase can further increase the versatility of this approach and allow highly efficient integration of genes surrounded by short homology arms into the chromosome in the absence of selection markers.

## Acknowledgements

We thank Dr. David Leach for insightful discussions, Dr. Aleksei Agapov for help with figure preparation.

## Funding

Russian Science Foundation grant 19-14-00359 (DNA interference assays).

Russian Ministry of Science and Higher Education grant 075-15-2021-1062 (recombination assays).

## Author contributions

Conceptualization: DE, AA and AK

Supervision: DE, AK

Investigation: DE, A. Okhtienko, A. Olina, MP

Data analysis: A. Okhtienko, MP, AK, DE

Preparation of figures: A. Okhtienko, A. Olina

Writing – original draft: AK, DE

Writing – review & editing: all the authors

## Competing interests

Authors declare that they have no competing interests.

## Data and materials availability

All data are available from the corresponding authors upon request. Small DNA libraries are available from the Gene Expression Omnibus (GEO) database. The code used for data analysis is available at the GitHub repository at https://github.com/Aokht17/pAgo_Genome_editors.

## Supplementary Materials

### Materials and Methods

### Plasmids and strains

Plasmids and strains used in this study are listed in Supplementary Tables S3 and S4. *E. coli* cultures were cultivated in LB Miller broth (2% tryptone, 0.5% yeast extract, 1% NaCl) with the addition of appropriate antibiotics (ampicillin 200 μg/ml, kanamycin 50 μg/ml, spectinomycin 50 μg/ml, chloramphenicol 25 μg/ml), 1.8% agar was added for plates preparation. NEB Turbo strain was used for routine cloning. MG1655 and DE160 strains were used for most experiments.

For the construction of the pBAD_CbAgo_lacI plasmids containing *lacI* fragments of different lengths, the expression vector pBAD_CbAgo (*19*) was digested with SphI in the presence of Shrimp Alkaline Phosphatase (rSAP, NEB). LacI fragments were PCR amplified from the pET28 plasmid using primers containing SphI cut sites on both ends, then digested with SphI, gel-purified and cloned into the pBAD_CbAgo vector using T4 ligase. Colony PCR was used to pick up clones with insertions in the correct orientation. The cloned gene contained no promoter region. Decoding of the *lacI* gene was performed manually by making substitutions in its every 6^th^-8^th^ position, without changing the protein sequence, and with preferable choosing of codons with the same frequency as in wild-type gene. Decoded *lacI* was synthesized as an IDT gBlock and cloned into pBAD_CbAgo backbone as described above. pBAD_CS_yffP plasmids were obtained using the Gibson assembly reaction from PCR products corresponding to the pBAD backbone without genomic DNA homology regions (*araC, araBAD* promotor, *rrn* terminators), and to the 300 bp *yffP* gene or an 1000 bp *yffN-yffO-yffP* operon fragment amplified from the genomic DNA of DE160. The I-SceI cut site was introduced in primers used for PCR amplification. A mutated variant of the site with two nucleotide substitutions, which is cleaved less efficiently (T<u>T</u>GGGATAACAGGGTAA<u>A</u>) (*31*), was used to avoid complete plasmid degradation. Plasmids pBAD_lacI and pBAD_CbAgoYK_lacI (encoding CbAgo with substitutions Y472A, K476A in the MID pocket) were obtained by PCR cloning with overlapping primers using the pBAD_CbAgo_lacI as the template. pBAD_CbAgoCD_lacI was obtained by excision of the catalytically dead CbAgo gene (with substitutions D541A, D611A) from pBAD_AgoCD (*19*) using NcoI and EcoRI and cloning it into pBAD_CbAgo_lacI treated with the same enzymes. The pDE351 plasmid used in recombination assays was obtained using the Gibson assembly reaction with HiFi mastermix (NEB) from 5 PCR products: (1) pSC101 *ori* from the pKD46 plasmid; (2) Sp^R^ gene from the pSyn6 vector (GeneArt, Thermofisher); (3) 4144 bp left homology arm including the *mhpR, mhpA, mhpB, mhpC* genes amplified from genomic DNA of MG1655; (4) the *cat* gene; (5) 5987 bp right homology arm including *cynX, lacA, lacY, lacZ* genes amplified from genomic DNA of MG1655.

The DE160 strain was obtained after transfer of the Z1 cassette with an inserted CbAgo gene under control of the tetP promoter from the donor strain DE157 (*19*) to recipient strain DL2988 containing I-SceI under control of the araBAD promoter, using P1 transduction.

#### CbAgo expression and purification

Plasmids from the pBAD_CbAgo series with inserted *lacI* fragments were transformed into *E. coli* MG1655 and incubated overnight at 37 °C on ampicillin LB plates. On the next day, all cells were inoculated into 1 L of LB media supplemented with 0.1% L-arabinose and ampicillin and grown for 14-20 h at 18 °C until OD_600_ = 1. The cells were collected by centrifugation (6000 g, 6 min at 4 °C) and pellets were frozen at -20 °C. Plasmids from the pBAD_CS_yffP series were transformed into the DE160 strain and the cells were grown in the same conditions with the addition of anhydrotetracylne in liquid media for CbAgo induction. Cell pellets were defrosted at room temperature and resuspended in 30 ml of lysis buffer (40 mM Tris-HCl pH 7.9, 150 mM NaCl) in the presence of protease inhibitor cocktail EDTA-free (Roche). After filtration using 170 µm nylon filters, the samples were lysed using a high-pressure homogenizer Cell Disruptor PEC at 30 kpsi twice and then centrifuged (17000 rpm, R21A Hitachi rotor, twice for 15 min at 4 °C). Clarified lysate was incubated with 0.3 ml of prewashed Co^2+^ beads (Clontech) for 140 min at +4 °C with rotation. The suspension was centrifuged for 3 min at 500 g at +4 °C, and supernatant was removed. Beads were washed twice with 25 ml of ice-cold lysis buffer for 5 min, 4 times with 1 ml of lysis buffer cootaining 10 mM imidazole, and CbAgo was eluted with lysis buffer containing 200 mM imidazole (3 times with 0.33 ml). 0.3 ml of each elution fraction was treated with Proteinase K (100 μg, 1 h 37 °C), and the fractions were combined for further gDNA purification.

#### Preparation of gDNA libraries

Deproteinized samples were treated with 0.5 ml of phenol-chloroform-isoamyl alcohol (25:24:1) pH 7.5-8, vortexed for 10 seconds, and centrifuged for 2 min at 21000 g. The upper aqueous phase was treated twice with 0.5 ml of chloroform. Nucleic acids from the final aqueous phase were ethanol-precipitated in the presence of PINK coprecipitant and 30 mM sodium acetate pH 5.0 for 1h in liquid nitrogen or overnight at -70 °C. Samples were centrifuged for 30 min, 21000 g at +4 °C and the pellets were washed with 70% of ice-cold ethanol. Air-dried pellets were dissolved in nuclease-free water (40 µl per sample). 10% of each sample was dephosphorylated using rSAP (NEB) in 1x T4 polynucleotide kinase (PNK) buffer for 30 min at 37 °C, rSAP was inactivated for 10 min at 75 °C. Dephosphorylated samples were radiolabelled with γ-P^32^-ATP using T4 PNK (NEB), an oligonucleotide ladder (12-70 nt) was also radiolabelled at this point. The labeled samples were mixed with the rest of corresponding untreated samples (20 µl), resolved by 19% PAGE with 8M urea in 1xTBE, and visualized using a Typhoon FLA 9500 imager (GE Healthcare). Gel slices containing 14–22 nt smalll DNAs were cut from gel, crushed in 0.4M NaCl and incubatted overnight at 20_°C with constant shaking (1000 rpm) on a bench tube shaker. Gel slices were removed and nucleic acids were ethano-precipitated as described above. DNA was dissolved in 20 μl of nuclease-free water. Small DNAs were ligated with Illumina-compatible adaptors 5’-adaptor -5’-GTTCAGAGTTCTACAGTCCGACGATC; 3’-linker - /5’P/TGGAATTCTCGGGTGCCAAGGAACTC/3’ddC/ bridge 1 - /5’AmMC6/CACCCGAGAATTCCANNNNNN/3’AmMO/ bridge 2 - /5’AmMC6/NNNNNNGATCGTCGGACTGTA/3’AmMO/).

The samples (20 μl each) were mixed with 8 μl of 5x Rapid Ligation buffer (ThermoFisher), 2 μl of 100 μM 5′-adaptor, 2 μl of 100 μM of 3′-linker, 2 μl of 100 μM bridge 1, 2 μl of 100 μM bridge 2 and 800 units of T4-ligase (NEB) and incubated for 16 h at room temperature. Ligated DNA fragments were sliced from 19% urea PAGE and eluted from the gel as described above. The libraries were PCR-amplified with RP1 and indexing primers (TrueSeq) using the NEBNext Ultra II Q5 Master Mix with adjusting the number of cycles after preliminary product visualization on 6% native PAGE with SYBR Gold staining. Amplified libraries wereseparated by native 6% PAGE (using visualization in blue light), extracted in 0.4 M NaCl and precipitated as described above. Small DNA libraries were sequenced using the HiSeq2500 platform (Illumina) in the rapid run mode (50-nucleotide single-end reads).

#### Small DNA analysis

All libraries were quality checked with FastQC (v0.11.9). Trimmomatik (v0.36) was used used to remove adapters, eliminate reads shorter than 14 bp and cut reads longer than 24 bp. Reads were aligned onto the reference genome of *E. coli* (MG1655 Refseq: NC_000913.3 or BW25113 Refseq: CP009273.1 with manually added CbAgo gene (802798-810180) and plasmid (pBAD_CbAgo_lacI_full, pBAD_CbAgo_lacI_dec or pBAD_yffP1000_wtCS) via bowtie (v1.3.1) allowing zero mismatches and unique alignment (-v 0 -m 1). Potential multi-mappers failed to align with the -m option were realigned using options: -a --best --strata -v 0 -m 10000. Multi-mappers that were aligned to both genome and plasmid sequences were not included in further analysis to avoid biases in coverage. The remaining reads were realigned with the same parameters (bowtie -a --best --strata -v 0 -m 10000). Uniquely aligned reads and selected multi-mappers were sorted and combined via samtools (v1.15). The number of reads for each multi-mapper was divided by the number of aligned sites. The *E. coli* genome was divided into 1 kb intervals using a custom Python script. To obtain read coverage within each interval, the bedtools (v2.30.0) was used to intersect the resulted bed file with the bam file. Small DNA coverage was expressed as RPKM (reads per kilobase per million aligned reads in the library). Reads from the plus and minus genomic strands were selected with samtools view –F/ -f 16; the coverage was calculated separately for each strand as described above. To find Chi sites at a given interval of the genome, a Python script was used that searches for the Chi sequences (5’-GCTGGTGG for the plus strand and 5’-CCACCAGC for the minus strand) within given coordinates. To calculate the fraction of smDNAs generated at the sites of DNA interference, small DNAs mapped to the chromosomal region between by the second closest Chi sites around the locus of interest were divided by the total number of chromosomal small DNAs in the library.

#### Analysis of the expression of the target loci during DNA interference

The same *E. coli* cultures that were used to obtain libraries of CbAgo-associated small DNAs were used to purify RNA by GeneJET RNA Purification Kit (ThermoFisher). RNA was treated with RNAse free DNAse (Qiagen). 2 μg of RNA was used in reverse transcription reaction with RevertAid reverse transcriptase and random hexamer oligonucleotide (ThermoFisher). The resulting cDNA was used in qPCR reactions with oligonucleotides specific for the genes of interest (*lacI, lacZ*, and *araC*) and the housekeeping gene *gapA*, using qPCRmix-HS SYBR premix (Evrogen) on a C1000 Touch Cycler with CFX96 Opticle Reaction Module (BioRad). Oligonucleotides for qPCR (synthetized by Evrogen) were selected by their specificity and efficiency, validated by Primer-BLAST and Multiple Primer Analyzer (ThermoFisher), PCR product melt curve and PCR efficiency evaluation, performed in serial dilutions experiments. Oligonucleotides were selected in such a way that they are active only with genomic cDNA roducts and cannot amplify products obtained from the plasmid template, which was additionally validated. For each reaction condition, three biological replicates were performed, each with three technical replicates. Oligonucleotides used for qPCR: lacI-for CGCTCACAATTCCACACAAC, lacI-rev CCGTCTCACTGGTGAAAAGAA, lacZ-for TATTGGCTTCATCCACCACA, lacZ-rev GTGCGGATTGAAAATGGTCT, araC-for AAAAAATGCACCGGGGCCAG, araC-rev CGGGTAGAATCAAACCGACCAG, gapA-for CTGGTGCGAAGAAAGTGGTT, gapA-rev GTCCTGGCCAGCATATTTGT.

#### *In vivo* recombination assay

Chemically competent cells of MG1655 were co-transformed with two plasmids (pDE351 and variants of pBAD_CbAgo). Some double transformants were obtained via two-step transformation (first with pBAD_CbAgo and second with pDE351). Co-transformed cultures were plated on LB agar plates supplemented with ampicillin, spectinomycin, chloramphenicol and D-glucose (1%), to prevent premature CbAgo expression. Individual colonies were inoculated in 6 ml of LB containing Amp+Chl+Sp+Glc and grown overnight. 1 ml of the cultures was mixed with 1 ml of 50% sterile glycerol and 100-200 μl aliquotes were frozen in liquid nitrogen and stored at -70 °C to use as start cultures in further experiments. Another 5 ml were used for miniprep plasmid DNA purification using Zymo Research D4020 kit. To confirm the presence of both plasmids, the samples were treated with specific restriction endonucleases Bsp1407I (linearizes pDE351) and BamHI (linearizes pBAD_CbAgo and its variants) and the products were analyzed by agarose gel electrophoresis.

For the recombination experiments, the starting cultures from -70 °C were defrosted at room temperature and 5 μl were inoculated into 1 ml LB (in 25 ml tubes) supplemented with 0.1% L-arabinose, ampicillin and chloramphenicol, or 1% Glc, ampicillin and chloramphenicol in control experiments, and grown at 30 °C for 24 hours. 10 μl of cells were washed with 0.3 ml LB+Glc to remove arabinose and prevent further CbAgo expression, inoculated into 0.5 ml LB+Glc in a 1.5 ml safe-lock tube and incubated overnight at 43 °C to induce loss of the pDE351 plasmid. This procedure was repeated 6 times in total (∼12 hours each), using 1 μl of cell cultures from the previous passage. Cells from the last 6^th^ passage were used for CFU counting on LB, LB+Cm, LB+Sp, and LB+Cm+Sp plates. Eight serial 10-fold dilutions were prepared in LB media for each culture in 96 well plates and then 10 μl of each dilution was placed on all plates. The plates were air-dried and incubated overnight at 37 °C, followed by colony counting.

**Fig. S1.**
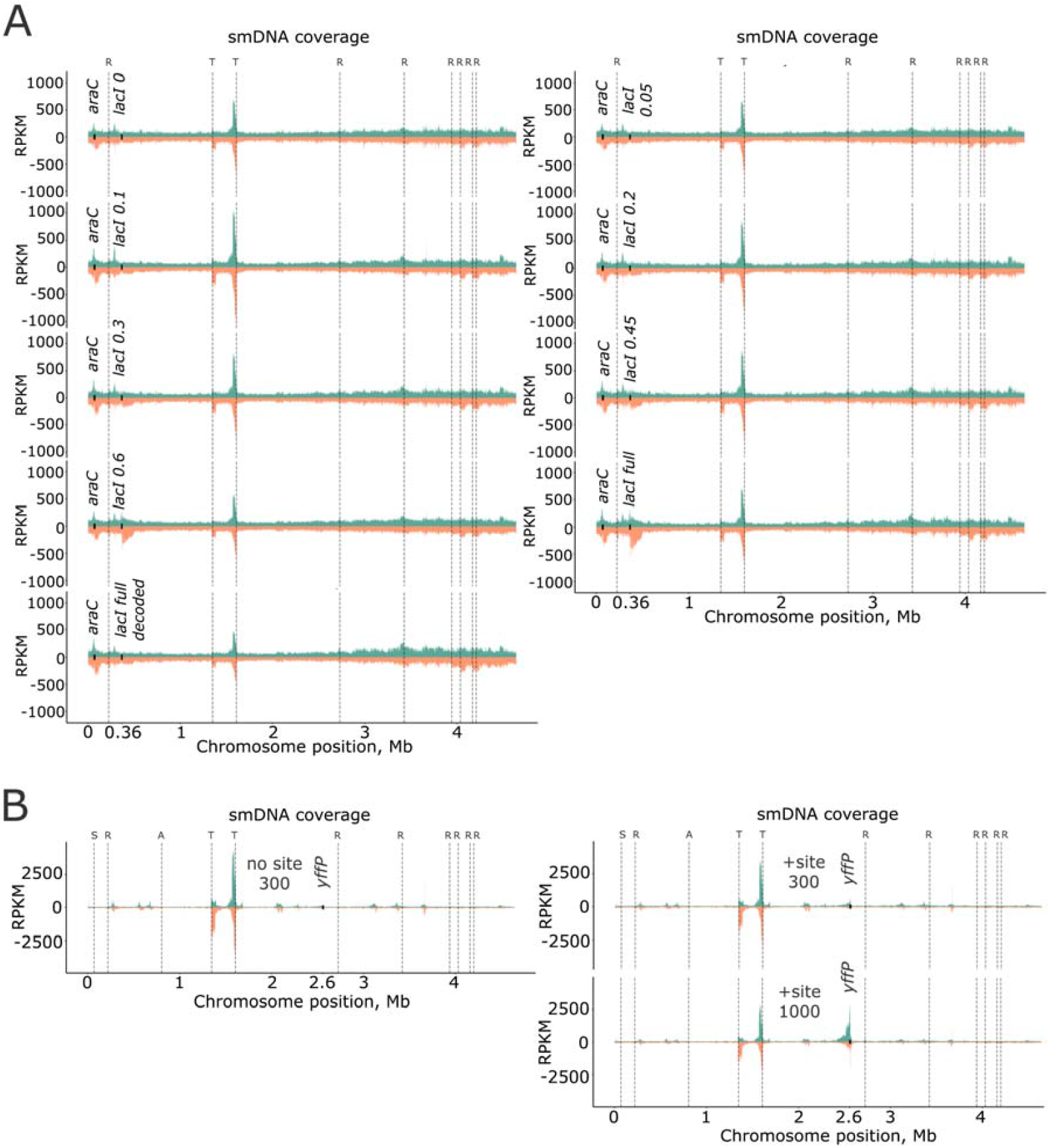
Whole-genome profiles of gDNAs associated with CbAgo in *E. coli* cells. (A) Genomic distribution of gDNAs in *E. coli* MG1655 containing pBAD_CbAgo_lacI plasmids with fragments of the *lacI* gene. (B) Genomic distribution of gDNAs in *E. coli* strain DE160 containing pBAD_yffP plasmids with or without the I-SceI cut site and 300 bp or 1000 bp region of homology from the *yffP* operon. The amounts of gDNAs are shown in reads per kilobase of genomic DNA per million reads (RPKM). Genes of interest are indicated (*araC, lacI, yffP*), positions of *terA* and *terC* sites (T), ribosomal RNA operons (R), I-SceI gene (S), CbAgo gene (A) are indicated.

**Fig. S2.**
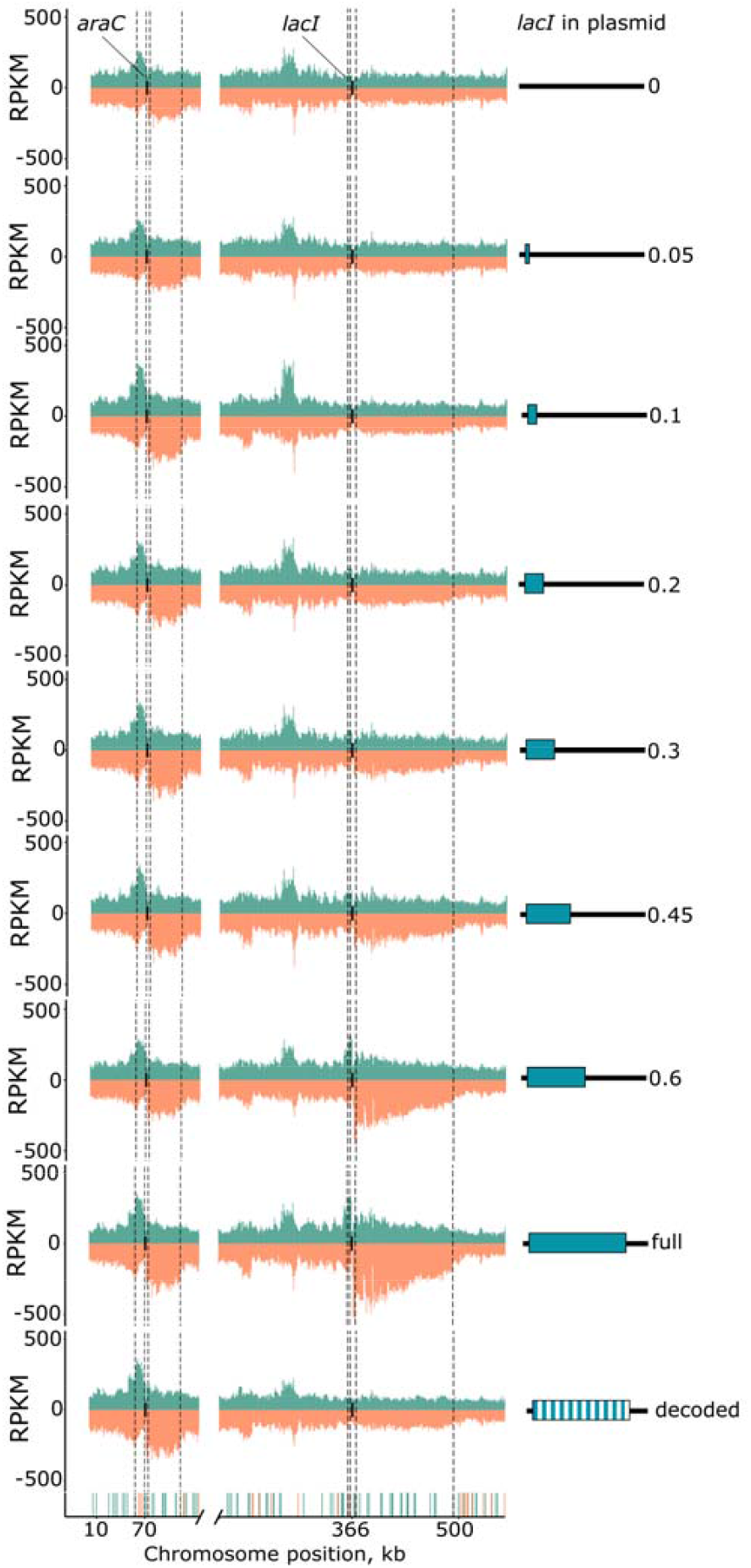
Analysis of genomic coverage of CbAgo-associated gDNAs in the *araC* and *lacI* loci in strains containing plasmids with fragments of the *lacI* gene. Strand-specific distribution of gDNAs around the *araC* (left) and *lacI* (right) genes is shown for each gDNA library; reads from the plus and minus genomic strands are shown in green and orange, respectively. Boxes in green correspond to the length of the *lacI* fragment on the plasmid. The closest Chi sites surrounding the target genes in the proper orientation are indicated (forward for the plus strand and reverse for the minus strand).

**Fig. S3.**
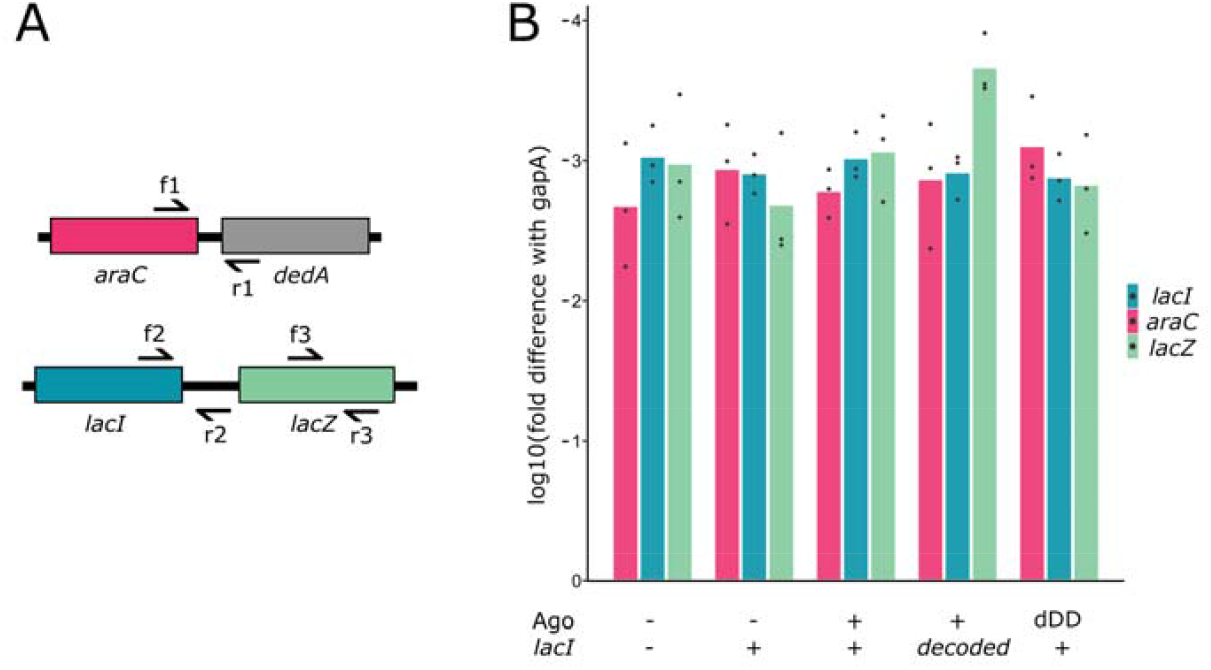
Analysis of the transcription levels of target chromosomal genes during CbAgo-induced DNA interference. (A) Scheme of chromosome-specific primers used for analysis of *lacI* and *araC* expression. Note that the plasmid *lacI* gene lacks promoter region but it still might be transcribed due to readthrough transcription from outside promoters. To avoid detection of plasmid transcripts resulting from background transcription of the *lacI* and *araC* genes, chromosome-specific primers were used for cDNA synthesis (r1, r2 or r3). Indicated pairs of primers were used for qPCR analysis of each gene. (B) Levels of *lacI* and *araC* expression relative to the *gapA* control. Means and standard deviations from three independent biological replicates are shown (each biological replicate is the average of three technical replicates performed with the same cDNA sample).

**Table S1.** Analysis of the ratio of gDNAs mapped to the plasmid and chromosomal sequences in experiments from Fig. 1.

| gDNA library | mapped reads, total | plasmid reads | plasmid to genome ratio, observed/expected<sup>a</sup> |
| --- | --- | --- | --- |
| 0 *lacI* | 2791323 | 179331 | 3.4 |
| 0.05 *lacI* | 4944884 | 306516 | 3.3 |
| 0.1 *lacI* | 3764124 | 327175 | 4.6 |
| 0.2 *lacI* | 4012629 | 294456 | 3.9 |
| 0.3 *lacI* | 3441276 | 272799 | 4.2 |
| 0.45 *lacI* | 2821147 | 211025 | 4.0 |
| 0.6 *lacI* | 2516040 | 228215 | 4.8 |
| full *lacI* | 5098616 | 511176 | 5.3 |
| decoded *lacI* | 2258945 | 375874 | 8.8 |

<sup>a</sup> The expected ratio was calculated based on the relative length of the replicons (4641652 bp for chromosomal DNA and 7421 bp for the plasmid containing full-length *lacI*) and the known copy number of the pBAD plasmid (12 copies on average).

**Table S2.** Percentage of *E. coli* cells resistant to chloramphenicol, spectinomycin and to both antibiotics in bacterial cultures in experiments on CbAgo-induced recombination between the editing plasmid and the chromosome. The percentage of resistant cells was calculated after plating of serial dilutions of all the cultures in the absence of antibiotics and in the presence of Cm, Sp, or Cm and Sp, and by dividing the numbers of resistant cells to the total CFU numbers in each culture. The data from 13-14 indeendent biological replicates are sorted in the descending order. Median values (or the closest data ponts) are indicated in bold.

| no CbAgo,<br>+Glc, +*lacI* |  |  | WT CbAgo,<br>+Glc, +*lacI* |  |  | WT CbAgo,<br>+Ara, -*lacI* |  |  | WT CbAgo,<br>+Ara, +*lacI* |  |  | dDD CbAgo,<br>+Ara, +*lacI* |  |  | dYK CbAgo,<br>+Ara, +*lacI* |  |  |
| --- | --- | --- | --- | --- | --- | --- | --- | --- | --- | --- | --- | --- | --- | --- | --- | --- | --- |
| Cm<sup>R</sup> | Sp<sup>R</sup> | Cm<sup>R</sup>Sp<sup>R</sup> | Cm<sup>R</sup> | Sp<sup>R</sup> | Cm<sup>R</sup>Sp<sup>R</sup> | Cm<sup>R</sup> | Sp<sup>R</sup> | Cm<sup>R</sup>Sp<sup>R</sup> | Cm<sup>R</sup> | Sp<sup>R</sup> | Cm<sup>R</sup>Sp<sup>R</sup> | Cm<sup>R</sup> | Sp<sup>R</sup> | Cm<sup>R</sup>Sp<sup>R</sup> | Cm<sup>R</sup> | Sp<sup>R</sup> | Cm<sup>R</sup>Sp<sup>R</sup> |
| 3.1 | 3.2 | 2.8 | 12.4 | 6.8 | 6.8 | 18.0 | 2.4 | 2.5 | 76.7 | 0.6 | 0.6 | 53.3 | 8.3 | 12.9 | 15.7 | 8.6 | 12.9 |
| 2.8 | 2.3 | 2.3 | 4.0 | 4.4 | 4.4 | 16.1 | 2.4 | 2.0 | 49.2 | 0.5 | 0.5 | 22.6 | 5.2 | 5.5 | 11.5 | 6.0 | 11.1 |
| 1.9 | 1.6 | 1.3 | 2.8 | 3.4 | 3.6 | 16.0 | 1.4 | 1.9 | 41.7 | 0.5 | 0.4 | 18.2 | 4.2 | 4.6 | 9.0 | 4.9 | 6.8 |
| 1.8 | 1.5 | 1.2 | 2.8 | 2.1 | 2.9 | 15.0 | 1.4 | 1.9 | 28.3 | 0.5 | 0.4 | 13.7 | 4.0 | 4.4 | 8.7 | 4.8 | 4.6 |
| 1.7 | 1.1 | 1.1 | 2.6 | 1.7 | 1.6 | 11.8 | 1.2 | 1.7 | 26.2 | 0.5 | 0.4 | 12.0 | 2.9 | 4.3 | 8.0 | 4.7 | 4.5 |
| 1.6 | 1.0 | 1.0 | 2.1 | 1.5 | 1.5 | 11.2 | 1.0 | 1.6 | 20.0 | 0.3 | 0.3 | 11.9 | 2.7 | 2.9 | 5.6 | 4.5 | 4.4 |
| **1.3** | **0.8** | **0.9** | **2.1** | **1.4** | **1.5** | **10.7** | **1.0** | **0.9** | **17.5** | **0.3** | **0.3** | **11.7** | **2.7** | **2.4** | **4.9** | **4.4** | **4.1** |
| **1.1** | **0.7** | **0.8** | 1.9 | 1.2 | 1.2 | **10.6** | **0.8** | **0.7** | **16.1** | 0.3 | 0.2 | **7.2** | **2.6** | **2.0** | **4.1** | **3.8** | **4.0** |
| 1.0 | 0.7 | 0.8 | 1.8 | 1.2 | 1.0 | 10.5 | 0.7 | 0.5 | 15.9 | 0.3 | 0.2 | 7.1 | 1.9 | 1.7 | 3.9 | 3.0 | 3.7 |
| 1.0 | 0.6 | 0.8 | 1.8 | 1.2 | 1.0 | 10.5 | 0.5 | 0.4 | 13.6 | 0.2 | 0.2 | 6.5 | 1.7 | 1.5 | 3.8 | 2.5 | 3.7 |
| 0.7 | 0.6 | 0.7 | 1.8 | 0.8 | 1.0 | 9.0 | 0.4 | 0.3 | 8.8 | 0.1 | 0.1 | 5.9 | 1.7 | 1.4 | 3.8 | 2.2 | 2.8 |
| 0.6 | 0.5 | 0.7 | 1.7 | 0.8 | 0.7 | 8.4 | 0.4 | 0.2 | 7.8 | 0.1 | 0.1 | 5.0 | 1.4 | 1.3 | 3.5 | 2.2 | 2.7 |
| 0.6 | 0.4 | 0.5 | 1.5 | 0.7 | 0.6 | 6.7 | 0.4 | 0.2 | 4.5 | 0.1 | 0.03 | 4.8 | 1.4 | 1.1 | 3.1 | 2.0 | 2.3 |
| 0.5 | 0.3 | 0.4 |  |  |  | 4.0 | 0.3 | 0.1 | 3.9 |  |  | 4.7 | 1.1 | 0.8 | 2.6 | 1.6 | 2.0 |

**Table S3.** Plasmids used in this study.

| Name | Description | Source |
| --- | --- | --- |
| pBAD_CbAgo | AmpR; codon-optimized N-terminally 6His-tagged CbAgo cloned into pBAD/HisB vector under control of the *araBAD* promoter | (19) |
| pBAD_CbAgo_lacI0.05 | AmpR; first 50 bp of the *lacI* gene (+1+50) cloned at the SphI site in pBAD_CbAgo | This study |
| pBAD_CbAgo_lacI0.1 | AmpR; 100 bp fragment of *lacI* (+1+100) cloned at the SphI site in pBAD_CbAgo | This study |
| pBAD_CbAgo_lacI0.2 | AmpR; 200 bp fragment of *lacI* (+1+200) cloned at the SphI site in pBAD_CbAgo | This study |
| pBAD_CbAgo_lacI0.3 | AmpR; 300 bp fragment of *lacI* (+1+300) cloned at the SphI site in pBAD_CbAgo | This study |
| pBAD_CbAgo_lacI0.45 | AmpR; 450 bp fragment of *lacI* (+1+450) cloned at the SphI site in pBAD_CbAgo | This study |
| pBAD_CbAgo_lacI0.6 | AmpR; 600 bp fragment of *lacI* (+1+600) cloned at the SphI site in pBAD_CbAgo | This study |
| pBAD_CbAgo_lacIFull | AmpR; full-length *lacI* (1083 bp) cloned at the SphI site in pBAD_CbAgo | This study |
| pBAD_CbAgo_lacI_decoded | AmpR; decoded full-length *lacI* (1083 bp) cloned at the SphI site in pBAD_CbAgo | This study |
| pBAD_yffP300_noCS | AmpR, pBAD vector with removed genome homology regions (*araC*, *araBAD* promoter and *rrn* terminators), and cloned 300 bp fragment of the *yffP* gene | This study |
| pBAD_yffP300_mutCS | AmpR, pBAD vector with removed genome homology regions (*araC*, *araBAD* promoter and *rrn* terminators), cloned 300 bp fragment of the *yffP* gene, and engineered cut site of I-SceI | This study |
| pBAD_yffP1000_mutCS | AmpR, pBAD vector with removed genome homology regions (*araC*, *araBAD* promoter and *rrn* terminators), cloned 1000 bp fragment of the *yffP* gene, and engineered cut site of I-SceI | This study |
| pDE351 | SpRCmR; editing plasmid; pKD46-derived thermosensitive origin pSC101 + SpR gene | This |
|  | from the pSyn6 vector (GeneArt, Thermofisher) + 5 kb homology arms corresponding to upstream and downstream chromosomal DNA around the *lacI* gene, with the *cat* gene cloned instead of *lacI* cloned using HiFi Assembly mastermix (NEB E2621) | study |
| pBAD_*lacI*Full | AmpR; full-length *lacI* (1083 bp) cloned at the SphI site in pBAD/HisB | This study |
| pBAD_CbAgoCD_*lacI*Full | AmpR; full-length *lacI* (1083 bp) cloned at the SphI site in pBAD_CbAgoCD encoding catalytically dead CbAgo (D541A, D611A) | This study |
| pBAD_CbAgoYK_*lacI*Full | AmpR; full-length *lacI* (1083 bp) cloned at the SphI site in pBAD_CbAgoYK encoding CbAgo with substitutions in the guide-binding pocket (Y472A, K476A) | This study |

**Table S4.**
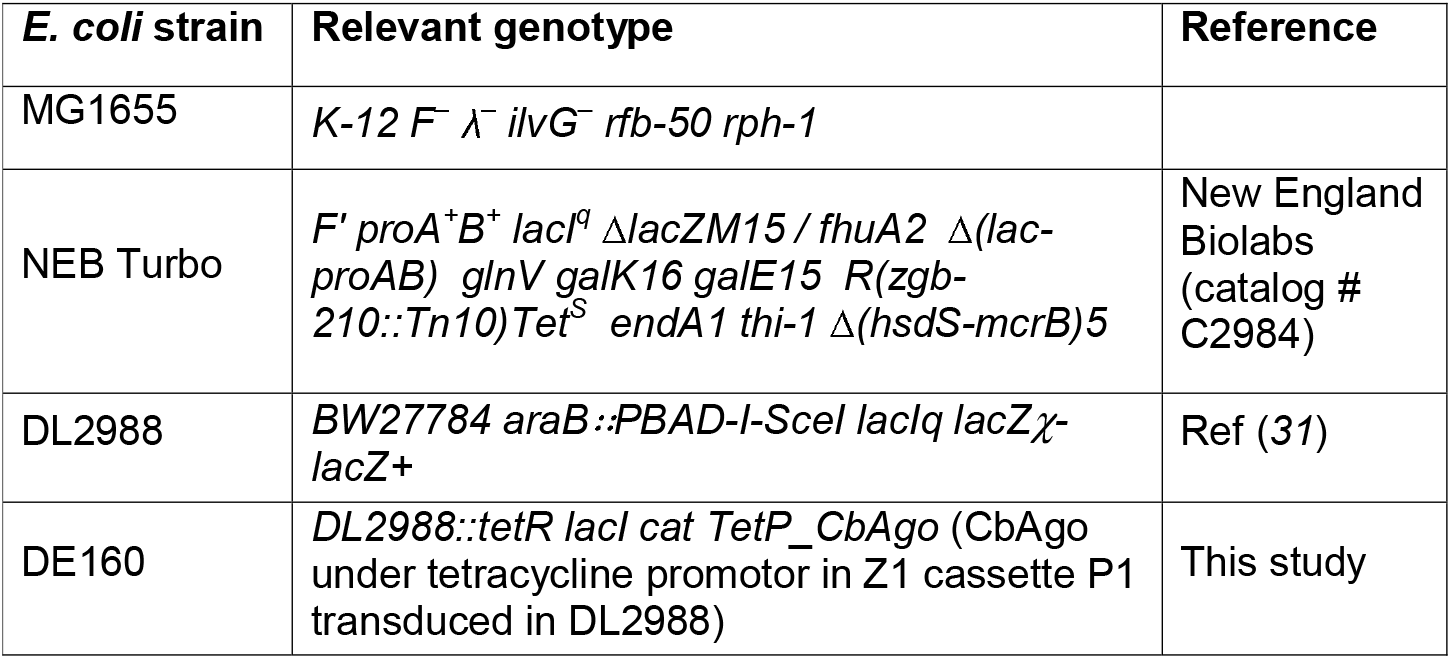
Bacterial Strains.

## Notes

### Competing Interest Statement

The authors have declared no competing interest.

